# Niclosamide shows strong antiviral activity in a human airway model of SARS-CoV-2 infection and a conserved potency against the UK B.1.1.7 and SA B.1.351 variant

**DOI:** 10.1101/2021.04.26.441457

**Authors:** Anne Weiss, Franck Touret, Cecile Baronti, Magali Gilles, Bruno Hoen, Antoine Nougairède, Xavier de Lamballerie, Morten Otto Alexander Sommer

## Abstract

SARS-CoV-2 variants are emerging with potential increased transmissibility highlighting the great unmet medical need for new therapies. Niclosamide is a potent anti-SARS-CoV-2 agent that has advanced in clinical development. We validate the potent antiviral efficacy of niclosamide in a SARS-CoV-2 human airway model. Furthermore, niclosamide is effective against the D614G, B.1.1.7 and B.1.351 variants. Our data further support the potent anti-SARS-CoV-2 properties of niclosamide and highlights its great potential as a therapeutic agent for COVID-19.

## Main Body

Since its emerge in 2019, coronavirus disease 2019 (COVID-19) caused by severe acute respiratory syndrome coronavirus 2 (SARS-CoV-2) led to over 3.1 million deaths worldwide as of April 26, 2021 (1). A tremendous joint research effort led to the approval of several vaccines at unprecedented speed yet anti-viral treatment options remain limited. At the same time, several viral variants harboring mutations in the N-terminal (NTD) and receptor-binding domain (RBD) of the spike protein gene, such as the B.1.1.7 (also named 20I/501Y.V1), B.1.351 (also named 20H/501Y.V2) variants, are causing global concern as they have been associated with enhanced transmissibility and possible resistance to vaccines and antibody neutralization (2–6). The B.1.1.7 and B.1.351 lineages have been linked to a ~50% increased transmission of SARS-CoV-2 infection and the vaccine efficacy of ChAdOx1 nCoV-19 has been reported to be reduced to 10.4% against the B.1.351 variant (6–9). Thus, despite the recent vaccine roll-out, there remains a high unmet need for novel therapeutics against SARS-CoV-2, which should be effective against circulating and potentially emerging variants of concern of SARS-CoV-2.

Niclosamide has been identified as a potent inhibitor of SARS-CoV-2 *in vitro* and *in vivo* and its optimized formulation for intranasal application and inhalation, was well-tolerated in healthy volunteers in a Phase 1 trial (10–13). Herein, we sought to further characterize the anti-viral properties of niclosamide by determining its potency in a human epithelial airway model of SARS-CoV-2 infection and tested its efficacy against several variants of concern of SARS-CoV-2.

To strengthen the existing data on the potent antiviral activity of niclosamide with a preclinical model resembling the human respiratory tract, we employed a trans-well bronchial human airway epithelium (HAE) model infected with SARS-CoV-2. HAE cultured at an airway-liquid interface has been extensively used as an *in vitro* physiological model mimicking the human mucociliary airway epithelium to validate the effectivity of antivirals on infections in conducting airways (14–16). The effect of niclosamide on the replication of SARS-CoV-2 in the HAE bronchial model (Eptihelix) was determined as previously described by Touret *et al*. (17) and Pizzorno *et al*. (14).

Briefly, human bronchial epithelial cells were apically infected with the European D614G strain of SARS-CoV-2 (BavPat1/2020; obtained from EVA-GLOBAL) at a MOI of 0.1 and cultivated in basolateral media that contained different concentrations of niclosamide (in duplicates) or no drug (virus control) for up to 4 days. Media was renewed daily containing fresh niclosamide. Remdesivir was used as experimental positive control and non-treated samples as negative control. On day 4, samples were collected at the apical side and the viral titer was estimated with a TCID_50_ assay. Then, cells were lysed, and the intracellular viral RNA was extracted and quantified by qRT-PCR. The viral inhibition was calculated with the infectious titers by normalizing the response, having the bottom value as 100% and top value as 0%. The IC_50_ was determined using logarithmic interpolation (Y=100/(1+10^((LogEC50-X)*HillSlope) in GraphPad Prims 7. Statistical analysis was performed using the Ordinary One-way Anova with Dunnett’s multiple comparisons test.

Niclosamide exhibited anti-SARS-CoV-2 activity by reducing the infectious titer and intracellular RNA levels in the HAE model in a dose-responsive manner. Niclosamide treatment with concentrations ≥ 1 μM significantly reduced the infectious titer to below the level of detection at Day 4 post-infection, yielding an IC50 of 0.96 μM (Fig. 1A and 1C). Furthermore, treatment with concentrations ≥ 1 μM of niclosamide significantly reduced the intracellular viral RNA level reaching a maximum effect of a 3-fold reduction on Day 4 (Fig. 1B). These data validate the substantial anti-SARS-CoV-2 effect of niclosamide in a reconstituted human airway model.

**Figure 1:**
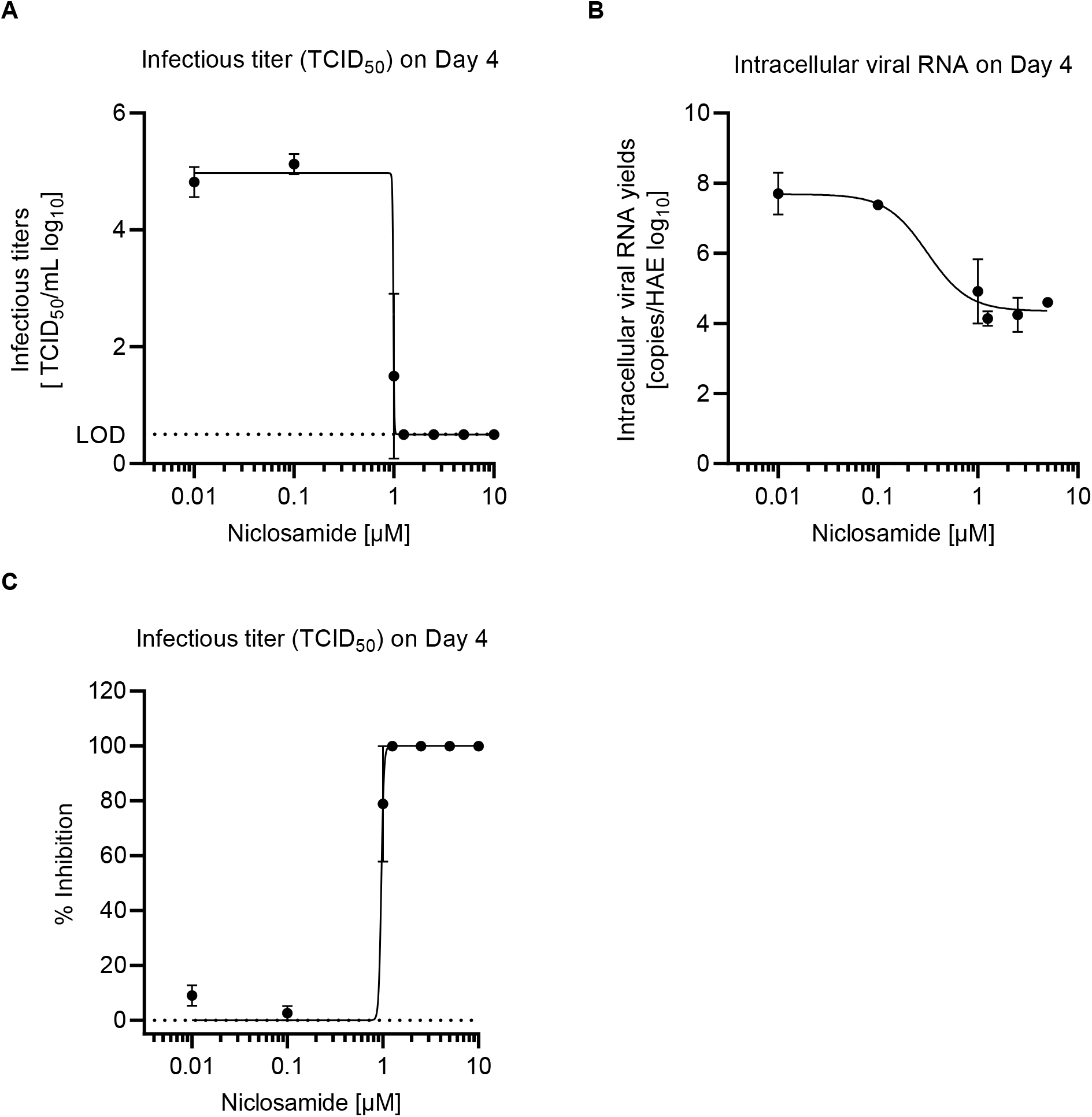
Antiviral efficacy of niclosamide in a trans-well model of human bronchial epithelium infected with SARS-CoV-2. Dose-dependent effects of niclosamide on infectious titer of SARS-CoV-2 (A) and intracellular viral RNA levels (B) on Day 4 post-infection. The reduction of infectious titer and intracellular RNA was significant for concentrations ≥ 1 μM niclosamide (infectious titer: 1 μM = p < 0.05, 1.25 – 10 μM = p < 0.0001; intracellular viral RNA: 1, 2.5, 5 μM = p < 0.01, 1.25 = p < 0.001 compared to non-treated control; Ordinary One way Anova with Dunnett’s multiple comparisons test). The IC50 based on the infectious titer on Day 4 was 0.96 μM (C). N = 2

We then tested the activity of niclosamide against several variants of concern of SARS-CoV-2, including the BavPat1 strain (D614G), SARS-CoV-2 201/501YV.1 (UVE/SARS-CoV-2/2021/FR/7b; lineage B.1.1.7, ex UK), SARS-CoV-2 Wuhan D614, and SARS CoV-2 SA lineage B.1.351 (UVE/SARS-CoV-2/2021/FR/1299-ex SA) in VeroE6 TMPRSS2 cells (ID 100978, CFAR). All viruses were obtained through EVA GLOBAL. The IC50 were determined by RT-qPCR as previously described by Touret *et al.* (18). Briefly, eight 2-fold serial dilutions of niclosamide in triplicate were added to the cells 15 min prior to viral infection and incubated for 2 days at 37°C. Remdesivir was used as experimental positive control and non-treated samples as negative control. The viral genome was quantified by real-time RT-qPCR from the cell supernatant (17). The IC50 was calculated as described above. All data associated with this study are present in the paper.

Niclosamide inhibited replication of the SARS-CoV-2 original strain (Wuhan D614) in VeroE6 TMPRSS2 cells with an IC_50_ of 0.13μM and IC_90_ of 0.16 μM which is in accordance with previous studies (10, 11). Importantly, niclosamide also blocked the replication of the European BavPat D614G, UK B.1.1.7 and SA B.1.351 variant with an IC_50_ of 0.06 μM, 0.08 μM and 0.07 μM, respectively (Fig. 2). Thus, niclosamide is effective against all tested variants of SARS-CoV-2 having a similar potency across the different strains compared to the original Wuhan D614 strain.

**Figure 2:**
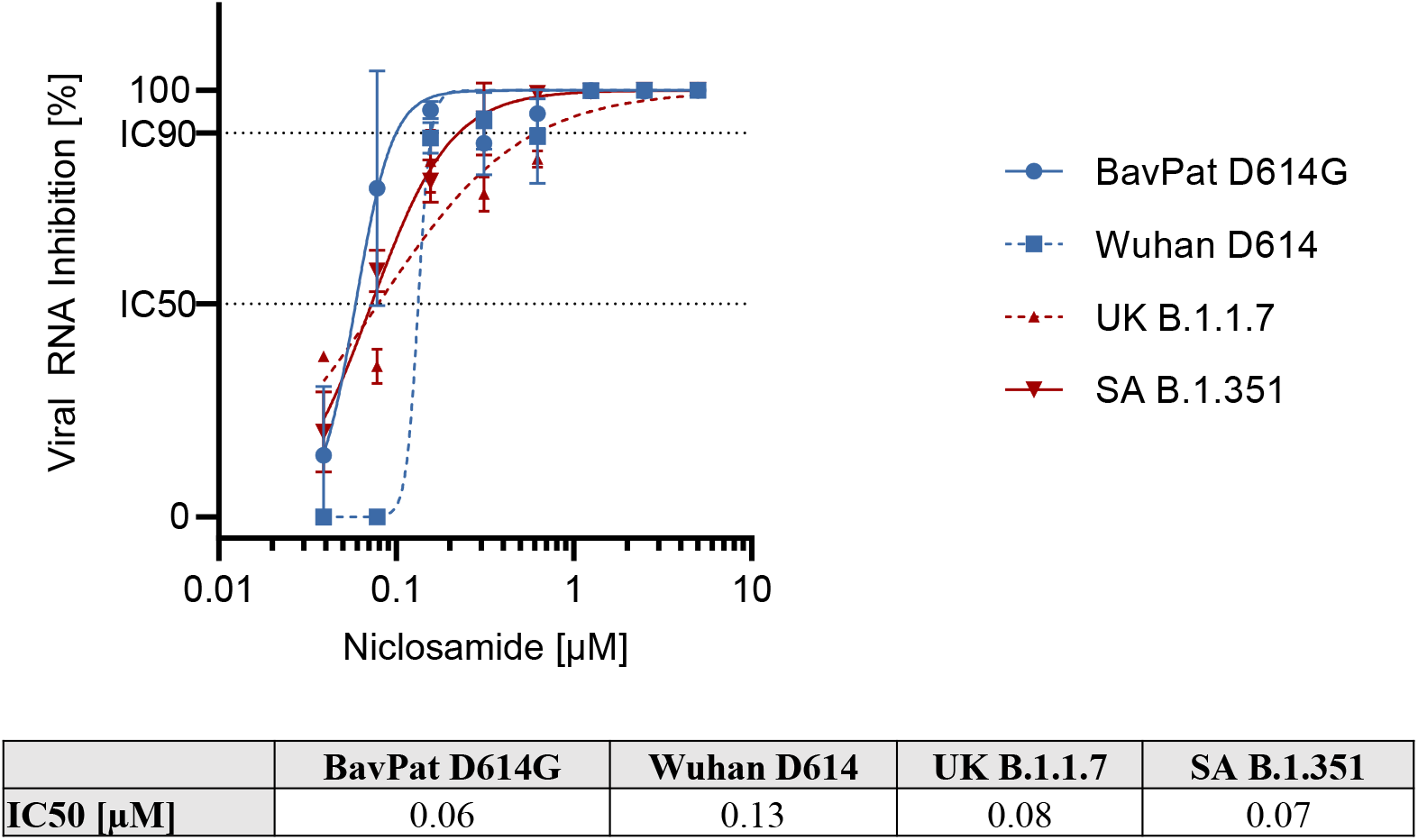
Effect of niclosamide on SARS-CoV-2 variants, including UK B.1.1.7 and SA B.1.351 in VeroE6 TMPRSS2 cells. IC = Inhibitory concentration. The origin of the tested variants is available at EVA-GLOBAL. N = 3

These data are in line with the host-targeted mode of action of niclosamide, which has been described to interfere with basic cellular mechanisms involved in SARS-CoV-2 replication, such as autophagy, the endosomal pathway and the TMEM16A chloride channel (11, 19–21). Accordingly, niclosamide is a potent antiviral therapeutic agent against SARS-CoV-2 and its variants. The molecule will also deserve further investigations to assess its potential role in the chemotherapeutic armamentarium required for future emerging infectious disease preparedness.

Taken together, our findings support niclosamide’s therapeutic potential as a potent anti-viral agent against SARS-CoV-2, including its variants of concern. Trials in patients with COVID-19 are needed to substantiate future clinical use.

## Acknowledgments

We are thankful for the support of Innovationsfonden Denmark (grant number: 0208-00081 and 0153-00209) and The Novo Nordisk Foundation under NFF grant number: NNF20CC0035580. We would like to thank Noemie Courtin (UVE) for her excellent technical assistance.

